# Chronic hM3Dq signaling in microglia ameliorates neuroinflammation in male mice

**DOI:** 10.1101/2020.01.27.921809

**Authors:** William Binning, Aja E. Hogan-Cann, Diana Yae Sakae, Matthew Maksoud, Valeriy Ostapchenko, Mohammed Al-Onaizi, Sara Matovic, Wei-Yang Lu, Marco A. M. Prado, Wataru Inoue, Vania F. Prado

**Affiliations:** Program in Neuroscience, University of Western Ontario, London, Ontario, Canada, N6A 5K8; Department of Physiology & Pharmacology, University of Western Ontario, London, Ontario, Canada, N6A 5K8; Department of Anatomy & Cell Biology, University of Western Ontario, London, Ontario, Canada, N6A 5K8; Robarts Research Institute, Schulich School of Medicine & Dentistry, University of Western Ontario, London, Ontario, Canada, N6A 5K8

**Author notes:** Corresponding authors: Vania F. Prado, Wataru Inoue, Marco A. M. Prado, Robarts Research Institute, The University of Western Ontario, 1151 Richmond St. N. London, ON N6A5B7. These authors contributed equally to this work (William Binning, Aja E. Hogan-Cann, Diana Yae Sakae). **Declarations of interest:** Nothing to declare.

**Keywords:** Cytokine, sickness behavior, microglia, muscarinic receptor, DREADD, GPCR, Gq

## Abstract

Microglia express muscarinic G protein-coupled receptors (GPCRs) that sense cholinergic activity and are activated by acetylcholine to potentially regulate microglial functions. Knowledge about how distinct types of muscarinic GPCR signaling regulate microglia function *in vitro* and *in vivo* is still poor, partly due to the fact that some of these receptors are also present in astrocytes and neurons. We generated mice expressing the hM3Dq Designer Receptor Exclusively Activated by Designer Drugs (DREADD) selectively in microglia to investigate the role of muscarinic M3Gq-linked signaling. We show that activation of hM3Dq using clozapine N-oxide (CNO) elevated intracellular calcium levels and increased phagocytosis of FluoSpheres *in vitro*. Acute treatment with CNO *in vivo* did not affect male mouse behavior, however chronic CNO treatment decreased sickness behavior triggered by lipopolysaccharide (LPS) treatment. Interestingly, whereas acute treatment with CNO increased synthesis of cytokine mRNA, chronic treatment attenuated LPS-induced cytokine mRNA changes in the brain, likely explaining the improvement in sickness behavior by chronic hM3Dq activation. No effect of CNO was observed in DREADD-negative mice. These results suggest that chronic activation of M3 muscarinic receptors (the hM3Dq progenitor) in microglia, and potentially other Gq-coupled GPCRs, preconditions microglia to decrease their response to further immunological challenges. Our results indicate that hM3Dq can be a useful tool to modulate neuroinflammation and study microglial immunological memory *in vivo*, which may be applicable for manipulations of neuroinflammation in neurodegenerative and psychiatric diseases.

**Highlights:**

- Microglial function was manipulated by specific activation of hM3Dq signaling.
- Chronic hM3Dq activation prevented LPS-induced sickness behavior in mice.
- Microglial hM3Dq signaling modulated expression of pro-inflammatory cytokines.

## 1. Introduction

Microglia are the resident immune cells of the central nervous system. They account for 5-10% of the total cells in the brain and are necessary for maintaining brain homeostasis (Li and Barres, 2018). Microglia are distributed throughout the CNS and dynamically survey the environment by extending and retracting their highly ramified processes. Upon environmental alterations, microglia quickly modify their activation state and adjust their range of effects which include phagocytosis of apoptotic cells, modulation of neuronal activity, elimination of synapses, production of neuromodulatory factors that support synaptic plasticity as well as release of cytokines to modulate inflammation in the adult brain (Kettenmann et al., 2011; Li and Barres, 2018; Wake et al., 2011).

It is increasingly recognized that microglia may be neurotoxic or exert beneficial roles depending on their activation status and release of mediators (Li and Barres, 2018). Among the multiple signals that operate on microglia to modulate their functional state and coordinate their diverse repertoire of functions are neurotransmitters. Microglia express numerous types of neurotransmitter receptors such as those for glutamate, GABA, serotonin, adrenaline and acetylcholine (Kettenmann et al., 2011; Li and Barres, 2018).

Cholinergic signaling is involved in the modulation of central and peripheral inflammation (Lehner et al., 2019; Pavlov and Tracey, 2015), however, we still have limited information about how microglia respond to activation of cholinergic receptors *in vivo*. Microglia have been reported to express the α3, α5, α6, α7 and β4 nicotinic channel subunits (Liu et al., 2009; Rock et al., 2008; Shytle et al., 2004; Suzuki et al., 2006). Expression of muscarinic receptors on cultured microglia has also been demonstrated (Pannell et al., 2016; Zhang et al., 1998). In cultured microglia, acetylcholine seems to exert mainly an anti-inflammatory effect on microglia. For instance, acetylcholine and nicotine inhibited TNF-α release in response to LPS stimulation (Shytle et al., 2004) and nicotine reduced free radical production in response to β-amyloid (Moon et al., 2008). How muscarinic receptors influence microglia function is poorly understood. On BV-2 microglia cells, acute activation of Gq-associated Designer Receptors Exclusively Activated by Designer Drugs (DREADDs) induced the production of pro-inflammatory mediators while activation of Gi-associated DREADDs inhibited lipopolysaccharide- and CCL2-induced inflammatory signaling (Grace et al., 2018). The influence of muscarinic receptors might be more important for the modulation of “activated microglia” than “resting microglia” as their expression is largely increased in microglia isolated from stroke and Alzheimer’s disease mouse models when compared to microglia isolated from healthy mice (Pannell et al., 2016). To note, most of these studies were done *in vitro* – and microglia have been shown to respond differently when analyzed *in vitro* or *in vivo* (Aguzzi et al., 2013; Bohlen et al., 2017; Yamasaki, 2014).

Here, we took advantage of DREADD technology to investigate the effects of M3 muscarinic Gq-mediated signaling on microglial function *in vivo*. We used the human modified muscarinic type 3 DREADD (hM3Dq), which is not activated by acetylcholine (Nichols and Roth, 2009) but can be activated by clozapine N-oxide (CNO) and its metabolites (Dong et al., 2010; Gomez et al., 2017), to explore the function of M3 muscarinic Gq-signaling *in vivo* in these cells. Using Cre/lox technology (Parkhurst et al., 2013) we generated a mouse line in which hM3Dq (Zhu et al., 2016) is selectively expressed in microglia. We found that acute activation of the hM3Dq pathway in microglia increased phagocytosis, increased the production of pro-inflammatory cytokines in the brain, but did not cause overt behavioral changes. In contrast, chronic activation of this signaling pathway attenuated the increase in brain cytokine production and sickness behavior triggered by low dose of lipopolysaccharide (LPS) injected peripherally. Our experiments provide a proof of principle that chronic M3 muscarinic Gq-linked signaling, and possibly other Gq-coupled GPCRs, can trigger immunological memory that helps to decrease inflammatory responses in the brain.

## 2. Materials and Methods

### 2.1. Animals

CX^3^CR1^CreER^ mice (Jackson Laboratory, B6.129P2(Cg)-Cx3cr1tm2.1(cre/ERT2)Litt/WganJ) expressing tamoxifen-activated Cre-recombinase in CX^3^CR1-positive cells (Parkhurst et al., 2013), were obtained from Jackson Laboratories. R26-hM3Dq/mCitrine mice (Zhu et al., 2016) (B6N;129-Tg(CAG-CHRM3*,-mCitrine)1Ute/J) expressing a CAG promotor-driven HA-tagged hM3Dq with an upstream floxed STOP cassette were a generous gift of Ute Hochgeschwender (Central Michigan University) and Bryan Roth (University of North Carolina). CX^3^CR1^CreER^ mice were crossed with R26-hM3Dq/mCitrine mice to create CX^3^CR1^CreER(+/-)^-hM3Dq(+/-) mice, which upon treatment with tamoxifen express hM3Dq in CX^3^CR1-positive cells (hM3Dq-positive mice). CX^3^CR1^CreER(+/-)^-hM3Dq(+/-) mice treated with vehicle (hM3Dq-negative mice) were used as controls. CX^3^CR1^CreER(+/-)^ or hM3Dq(+/-) littermate controls were used as additional hM3Dq-negative controls. To avoid expression of h3MDq in circulating myeloid-derived CX^3^CR1 positive cells, experiments were performed no sooner than 1 month following tamoxifen treatment (P90) to allow turnover of peripheral myeloid-derived cells, as previously described (Parkhurst et al., 2013).

All procedures were conducted in accordance with guidelines of the Canadian Council of Animal Care and approved by the University of Western Ontario Animal Use Subcommittee (protocol 2016-103 and 2016-104). The study also complied with the ARRIVE guidelines (Animal Research: Reporting *In Vivo* Experiments). Animals were housed maximum of four by cage (28×18 cm) and had access to rodent chow (Envigo 7913) and water *ad libitum.* They were maintained on a 12h/12h light-dark cycle in non-SPF conditions.

### 2.2. Experimental Design

Experimental groups consisted of 4-6 month old CNO-treated CX^3^CR1^CreER(+/-)^-hM3Dq(+/-) mice. Vehicle-treated CX^3^CR1*^CreE^*^R(+/-)^-hM3Dq(+/-) animals were used as controls. CX^3^CR1^CreER(+/-)^ littermate males were used as additional controls. Mice were randomly assigned to a treatment group and investigators were blinded during experimental procedures.

### 2.3. Drugs

Tamoxifen (Sigma, cat#T56648) was dissolved at 100 mg/ml in 10% ethanol/90% corn oil. To induce DREADD expression, tamoxifen (75 mg/kg; i.p.) was injected once per day for 5 consecutive days. For cell culture assays, the tamoxifen active metabolite 4-hydroxytamoxifen (4-OHT, Sigma, cat#H6278) was used instead of tamoxifen. 4-OHT was dissolved in 100% ethanol and used at concentration of 5 µM. Clozapine N-Oxide (CNO, Tocris, cat#4936) was dissolved in 0.5% dimethylsulfoxide (DMSO). Calcium dye Rhod-2 AM (Abcam, cat#ab142780) was used at a final concentration of 5 µM in 1% DMSO. Store-operated Ca^2+^ channel (SOCC) inhibitor SKF-96365 (Santa Cruz, cat#130495-35-1) was diluted in water and used at 20 µM. *E. coli* 0111:B4 lipopolysaccharide (LPS) (Sigma-Aldrich, cat#L3012, Ec#297-473-0, lot#WI150506) was dissolved in saline and used at a dose of 0.1 mg/kg, i.p. This dose of LPS was chosen because it is the minimum effective dose to elicit pro-inflammatory cytokine response and sickness behavior in mice (Berg et al., 2004).

### 2.4. Antibodies

Primary antibodies used include anti-HA (1:1000, BioLegend, cat#901501, RRID:AB_2563417), anti-Iba1 (1:1000, Abcam, cat#ab5076, RRID:AB_2224403), anti-GFAP (1:1000, Abcam, cat#ab7260, RRID:AB_296804), and anti-NeuN (1:1000, Cell Signaling, cat#D3S3I, RRID:AB_2630395). Secondary antibodies used include Alexa Fluor 555 goat anti-mouse (1:1000, ThermoFisher, cat#A32727, RRID: AB_2535783) and Alexa Fluor 633 goat anti-rabbit (1:1000, ThermoFisher, cat#A-21070, RRID:AB_10372583).

### 2.5. Immunohistochemistry

CX^3^CR1*^CreER^*-hM3Dq mice underwent intracardiac perfusion with 1X PBS for 5 min and 4% PFA in PBS for 5 min. Brains were extracted and postfixed in 4% PFA at 4°C for 24 h. Brains were then cut into 40-µm thick sections using a vibratome (VT1200S, Leica). Staining for Iba1, HA, NeuN or GFAP was performed as described previously (Kolisnyk et al., 2016) using Alexa Fluor 555/633-tagged secondary antibodies and Hoechst. Sections were mounted on glass slides using Fluorescent-G mounting medium (Electron Microscopy Sciences, cat#17984-25) and imaged using an SP8 confocal microscope (Leica) equipped with a 20x/0.75 dry or a 40x/1.3 oil objective. At least 4 sections were imaged for each animal (N=5) for each condition, with at least 6 fields imaged for each section. Images were analyzed using ImageJ Cell Counter plugin.

### 2.6. Primary microglia cultures

Mixed glia cultures were prepared from P0-P2 CX^3^CR1^CreER(+/-)^-hM3Dq(+/-) mice following an established procedure (McCarthy and de Vellis, 1980; Pinteaux et al., 2002) with small modifications. Briefly, extracted forebrains were dissociated in DMEM/F12 (Multicell, cat#319-075-CL) containing 10% heat-inactivated fetal bovine serum (Gibco, cat#10438026) and 1× penicillin/streptomycin (Gibco, cat#15140-122). Cells were plated on 75-cm^2^ tissue culture flasks coated with poly-D-lysine (10µg/mL, Sigma Aldrich, cat#P9155) and incubated at 37 °C in 5% CO_2_ atmosphere. The culture medium was changed after 3 days and then every 5 days until confluency (12–18 days *in vitro*).

To obtain pure microglia cultures, flasks of mixed glial cultures were shaken at 200 rpm for 2 h at 37 °C. Floating microglia were collected by centrifugation (380g for 10 min) and seeded at ∼1×10^5^ cells/ml on poly-D-lysine-coated dishes containing 50% old/50% new media. On the following day, culture medium was supplemented with 5 µM 4-OHT and incubated for 48 h for optimal induction of hM3Dq expression. Approximately 5×10^5^ cells were obtained per brain used.

### 2.7. Immunocytochemistry

Immunocytochemistry was done on microglia plated on poly-D-lysine-coated glass coverslips at a density of ∼1×10^4^ cells per well. Cells were obtained from three independent cultures of three different litters to confirm HA expression. Cells were fixed with 4% paraformaldehyde (PFA) in PBS. After permeabilization with 0.5% Triton X-100 and blocking with 5% bovine serum albumin (BSA)/0.1% Triton X-100, cultures were incubated with primary antibodies (anti-HA, anti-Iba1, and anti-GFAP) overnight at 4 °C. Cells were then washed with PBS and incubated for 1 hour with secondary antibodies (Alexa Fluor 555 goat anti-mouse and Alexa Fluor 633 goat anti-rabbit, 1:500) and Hoechst (1:2000). Coverslips were then mounted on glass slides using Immumount (Thermo Fisher) medium and imaged using an FV1000 confocal microscope (Olympus) equipped with a 20x/0.75, 30x/1.05 or 60x/1.35 objective. Images were analyzed using ImageJ (NIH, Bethesda, MD) Cell Counter plugin.

### 2.8. Calcium imaging

For Ca^2+^ imaging, microglia were plated on poly-D-lysine-coated glass-bottom dishes (MatTek). Ca^2+^ imaging was performed as described previously (Beraldo et al., 2016) with some modifications. Cultures were washed with Krebs-Ringer HEPES (KRH) buffer (129 mM NaCl, 5 mM NaHCO_3_, 4.8 mM KCl, 1.2 mM KH_2_PO_4_, 1.2 mM MgCl_2_, 2.8 mM glucose, 10 mM HEPES) supplied with 1 mM CaCl_2_, incubated for 45 min with 5 µM of red spectrum fluorescent dye Rhod-2 AM at 37 °C, washed again 3 times with calcium-free KRH and incubated in KRH with or without calcium for 10 min at 37 °C. Intracellular Ca^2+^ was recorded with a FV1000 confocal microscope equipped with a 30x/1.05 objective. The imaging field was chosen to contain cells expressing hM3Dq as shown by mCitrine fluorescence (ex488/em505-550). Time kinetics were done with a 2.5-second step. After 100 s of background recording, 3 µM CNO, 1 mM ATP and 2 µM ionomycin or 10 µM thapsigargin (for KRH with or without calcium, respectively) were added consecutively to the dish. Rhod-2 fluorescence was registered using 546 nm excitation laser and 560-660 nm emission filter. ROIs were assigned and quantified for each cell in the imaging field using ImageJ (NIH, Bethesda, MD). Integrated fluorescence was normalized to the average signal during background recording. At least three independent cultures were obtained from three different litters and at least 50 cells were imaged for each condition.

### 2.9. Voltage-clamp recordings

Control cells and 4-OHT-treated cells on poly-D-lysine-coated 35-mm petri dishes were kept in bath solution containing (in mM): 130 NaCl, 5 KCl, 3 MgCl_2_, 2 CaCl_2_, 10 HEPES and 5 glucose. For patch-clamp recordings, KCl in the bath solution was replaced with NaCl (containing 135 mM NaCl). Patch electrodes were pulled from thin-wall glass tubes (World Precision Instruments, Inc.; Sarasota, FL) using a PP-830 pipette puller (Narishige, Japan). The electrode was filled with intracellular solution containing (in mM): 140 Cs-gluconate, 8 NaCl, 1 MgCl_2_, 10 Cs-BAPTA, and 10 HEPES following a previously described protocol (Zweifach and Lewis, 1996) with modifications. The pH of all solutions was adjusted to 7.4 using NaOH and/or KOH, and the osmolarity was adjusted to ∼300 mOsm. Under voltage-clamp conditions, whole-cell patch recordings were performed using a Multiclamp 700B amplifier (Molecular Devices) by a person blinded to the treatment compounds. After measuring membrane capacitance of the test cell, voltage ramps (from −100mV to +50mV over 150 ms) were repeatedly applied to the cell every 10 s, revealing the current-voltage relationship. When stable recordings of baseline transmembrane current were achieved, the test cells were perfused with 3 µM CNO for 6 min, followed by application of 20 µM SKF-96365. The recorded electrical signal was digitized at 10kHz, low-pass at filtered (1 kHz), acquired on-line using the program pClamp (Molecular Devices), and saved in a computer for off-line analysis. The current density (pA/pF) at different voltages for each test cell was averaged and plotted for report. At least 15 cells from three different litters were analyzed for each testing condition.

### 2.10. Phagocytosis Assay

Microglia were plated on glass coverslips coated with poly-D-lysine (∼2×10^4^ cells/glass) and incubated in DMEM/F12 (Multicell, cat#319-075-CL) containing 10% heat-inactivated fetal bovine serum (Gibco, cat#10438026) and 1× penicillin/streptomycin (Gibco, cat#15140-122) for 1 day. After approximately 24 hours, 5 µM 4-OHT was added and incubated for another 2 days. Phagocytosis assay was performed as described previously (de Jong et al., 2008). Briefly, medium containing 0.05% (v/v) of 1.0 µm diameter blue fluorescent FluoSpheres carboxylate-modified microspheres (ThermoFisher, cat#F8814) was added to the wells for 45 min. Microglia cultures were then washed gently with ice cold PBS and fixed with 4% PFA for 20 min. Immunocytochemistry was then performed for HA and Iba1 as described above. Microglia cultures were prepared from three separate litters, both male and female pups were used for cultures. Number of cells, number of FluoSpheres, and number of cells containing FluoSpheres were quantified using ImageJ Cell Counter plugin.

### 2.11. RNA extraction and quantitative PCR

Total RNA was extracted from the hippocampus using Aurum Total RNA mini-kit (Bio-Rad, cat#732-6870) and phenol/chloroform mix (Invitrogen, cat#15593-031). cDNA was synthesized from 2 µg of RNA using High Capacity cDNA RT kit (Applied Biosystems, cat#4368814). qPCR was performed in a CFX96 Real-time system (Bio-Rad) using SensiFAST SYBR NO-ROX kit (Invitrogen, cat#BIO-98020) and specific primer sets (shown below) for TNF-α, IL-1β, IL-6, and RPL13a as the reference gene. Experimental groups consisted of at least 7 animals. Data were analyzed using the comparative threshold cycle (C_t_) method and results were expressed as fold of change from control. Specific oligonucleotide primers for qPCR are as follows: TNF-α, forward, 5’-CTTCTGTCTACTGAACTTCGGG-3’, reverse, 5’-CAGGCTTGTCACTCGAATTTTG-3’; IL-1β, forward, 5’-ACGGACCCCAAAAGATGAAG-3’, reverse, 5’-TTCTCCACAGCCACAATGAG-3’; IL-6, forward, 5’-GATGCTACCAAACTGGATATAATCAG-3’, reverse, 5’-CTCTGAAGGACTCTGGCTTTG-3’; and RPL13a, forward, 5’-CTGCCCCACAAGACCAAGAG-3’, reverse, 5’-GGACCACCATCCGCTTTTTC-3’.

### 2.12. Behavior

For all behavioral experiments, mice were injected with vehicle or CNO (1.0 mg/kg, i.p.) 30 min before the trial. All experiments were completed with 4-6 month old male CX^3^CR1^CreER(+/-)^- hM3Dq(+/-) mice during the light period (between 10:00 and 16:00). Three cohorts consisting of at least 8 mice per group were tested over three consecutive days. The first cohort was tested for open field and social preference. The second cohort was used for light-dark box, elevated plus maze, and forced swim test. The third cohort was used for social memory. Experimenter was blind to treatment.

#### 2.12.1. Open Field

Open field activity was assessed as previously described (Janickova et al., 2017; Martins-Silva et al., 2011) with some modification. VersaMax Tracking System (X-Y and Z sensors) automatically recorded activity of mice in an open field box (40.64×40.64 cm^2^) (Omnitech Electronics Inc., Columbus, USA) that was separated into 4 equal quadrants using acrylic plexiglass. Recordings were done without an investigator present in the room. Two mice were placed in each open field box, one in the lower left corner and one in the top right corner of the box. The task was completed on two consecutive days with the exact same criteria. Locomotor activity is expressed as the total distance (cm) that mice travelled during the trial, while time in center is measured as the total time (seconds) that mice spent within the inner (8×8 cm^2^) region of their quadrant. All open field data (locomotor activity and time in center) are expressed as a sum of the first 60 min of the trial. For all open field experiments assessing baseline activity, a repeated measures two-way ANOVA with Bonferroni post hoc analysis was used. For all open field experiments assessing LPS-induced neuroinflammatory behavior, student’s t-test was used for analysis.

#### 2.12.2. Light-Dark Box

Light-dark test was performed as previously described (Depino et al., 2008). Mice were automatically recorded with VersaMax Tracking System (X-Y and Z sensors) in an open field box (40.64×40.64 cm^2^) containing a dark enclosure (Opto-Varimex-5 Light/Dark Box, 0150-LDB) within the top half of the box. Mice were placed under the hole of the dark enclosure and allowed to freely move for 5 min. Data are represented as total time the mice spent in the half without the dark enclosure. The crossover measurement – which records time spent in one region until the animal’s body fully enters the other region – was used to prevent mice poking their heads out of the dark enclosure from being tracked as light side activity. Student’s t-test was used for analysis.

#### 2.12.3. Elevated Plus Maze

To assess anxiety-like behavior, the elevated plus maze was performed as previously described (Janickova et al., 2019). Additional details are as follows. A maze with two open arms and two enclosed arms (open arms: 30×5 cm, surrounded by a 0.5-cm-high border; closed arms: 30×5 cm, surrounded by 19-cm-high walls) (Med Associates Inc., St Albans, USA) was used. Mice were placed on the central platform facing an open arm and were allowed to freely explore for 5 min. ANY-maze tracking software (Stoelting Co.) recorded locomotor activity during the test. Total time mice spent in the open arms of the maze was compared using a Student’s t-test.

#### 2.12.4. Forced Swim Test

Forced swim test was completed similarly to a previously described procedure (Beraldo et al., 2015). A plastic beaker (diameter, 15 cm; height, 25 cm) filled with 16 cm of water at 25°C was placed in the center of a table underneath a camera with ANY-maze software. Mice were placed in the beaker of water, and immobility/mobility was recorded for 6 min. The first min of the trial was discarded to account for the habituation phase. Data expressed as time of immobility during the last 5 min of the trial were analyzed using Student’s t-test.

#### 2.12.5. Social preference

Sociability was assessed as previously described (Kwon et al., 2006). Briefly, testing was done in a three-chambered apparatus (15×90×18.5 cm divided into three chambers of 15×29 cm and separated by dividers with a central 3.8×3.8 cm door) (Med Associates, St. Albans, VT, USA) that offers the subject a choice between a social stimulus and an object. Mice were first habituated for 10 min to the three-chamber apparatus and two empty metal mesh cylinders at each end (Leite et al., 2016). Mice were then locked in the center of the chamber for 5 min, during which a novel object (green lego block) was placed under the cylinder at one end, and a juvenile mouse of different background was placed under the cylinder at the other end. During the 10 min task, time spent investigating the object and the juvenile mouse was defined as the time a mouse spent sniffing through the holes of either cylinder and was manually scored by an investigator blind to the treatment. In baseline experiments, data were analyzed using student’s t-test. In LPS experiments, data were analyzed using repeated measures two-way ANOVA with Bonferroni post hoc analysis.

#### 2.12.6. Social Memory

Social memory was completed as previously described (Prado et al., 2006). Briefly, mice were singly housed 4 days prior to the experiment to establish territorial dominance. Immediately before testing, mice were habituated to an empty metal mesh cylinder for 10 min in their home cage. To test social interaction, a juvenile intruder was then placed in the test subject’s home cage under the holed cylinder, allowing for olfactory investigation. Time of investigation was defined as the time a mouse spent sniffing through the holes of the cylinder and was manually recorded by an investigator blind to treatment. The task consisted of five trials, each 5 min long and separated with 15 min intertrial intervals. The same juvenile intruder was used for the first four trials and a new juvenile intruder was introduced in trial five to assess social memory. Time spent investigating the juvenile intruder during each of the 5 trials was compared using repeated measures two-way ANOVA with multiple comparisons with Bonferroni post hoc analysis.

#### 2.12.7. Object Recognition

In order to control for potential recognition deficits, an experimental design similar to social memory was used (Prado et al., 2006). Initially, a 10-min habituation phase was given where the mice explored an empty holed acrylic cylinder in the test cage. Mice then underwent five consecutive 3 min trials with 10 min intertrial intervals. A green rectangular lego block was placed under the cylinder in the first four trials, whereas a red rectangular lego block was used in the last trial. The time spent sniffing the acrylic cylinder (within 5cm; defined by Any-maze software) was measured in each of the five trials. Results are graphed as time (seconds) the mice spent investigating the cylinder during each trial and were analyzed using two-way ANOVA with repeated measures with Bonferroni post hoc analysis.

#### 2.12.8. Olfaction Test

Olfaction was assessed as previously described (Prado et al., 2006). Briefly, to test for olfactory deficits, mice were exposed to the same scent (vanilla extract) for four consecutive trials and were exposed to another scent (caramel extract) on the fifth trial. Each trial was 3 min in duration with 10 min intertrial intervals. A 5×5 cm^2^ paper towel was soaked with 3 large drops of either vanilla or caramel extract and placed in a small petri dish with several small holes in its cover to allow for olfaction but prevent physical interaction with the paper towel. The paper towel was replaced with a newly soaked one before each trial to maintain a consistent level of scent during each trial. Time of sniffing was manually recorded by an investigator blind to treatment. Time spent sniffing during each trial was compared using two-way ANOVA with multiple comparisons.

### 2.13. Statistical Analysis

All statistical analyses were performed using GraphPad software (Prism). For two-group comparisons, Student’s unpaired t-test was used when samples displayed a normal distribution. For multiple groups comparisons, one-way ANOVA was used followed by Tukey’s post hoc test when appropriate. For multiple factor comparisons, two-way ANOVA was used with Bonferroni’s post hoc test. Data are expressed as a mean ± standard error of the mean (SEM).

## 3. Results

### 3.1. hM3Dq is expressed selectively in Iba1-positive microglia in CX^3^CR1^CreER(+/-)^-hM3Dq(+/-) mouse brain

To examine how M3 muscarinic receptors influence microglia function we used the Cre-loxP strategy to generate mice expressing the human modified M3 muscarinic DREADD (hM3Dq) selectively in microglia. Specifically, we crossed the tamoxifen inducible Cre mouse line CX^3^CR1-CreER (Parkhurst et al., 2013) to the R26-hM3Dq/mCitrine mice (Zhu et al., 2016) to generate CX^3^CR1^CreER(+/-)^-hM3Dq(+/-), which express hM3Dq under control of the strong ubiquitous promoter CAG separated by the DREADD coding sequence by a floxed STOP cassette (Fig. 1A). The hM3Dq carries an HA-tag, which allows detailed spatial localization of the receptor in the cell and is bicistronic, as it is followed by a self-cleaving P2Apeptide and a mCitrine reporter (Zhu et al., 2016). Tamoxifen-induced Cre expression results in recombination (allowing removal of the STOP cassette) and persistent expression of hM3Dq in long-lived microglia. Conversely, short-lived myeloid cells (monocytes and inflammatory macrophages), which are rapidly replenished by CX^3^CR1*^-^* bone marrow precursors, do not carry Cre-mediated recombination 30 days after tamoxifen injection (Parkhurst et al., 2013). Thus, to permit Cre-dependent expression of hM3Dq almost exclusively in microglia, all *in vivo* experiments were conducted at least one month after tamoxifen treatment.

**Figure 1.**
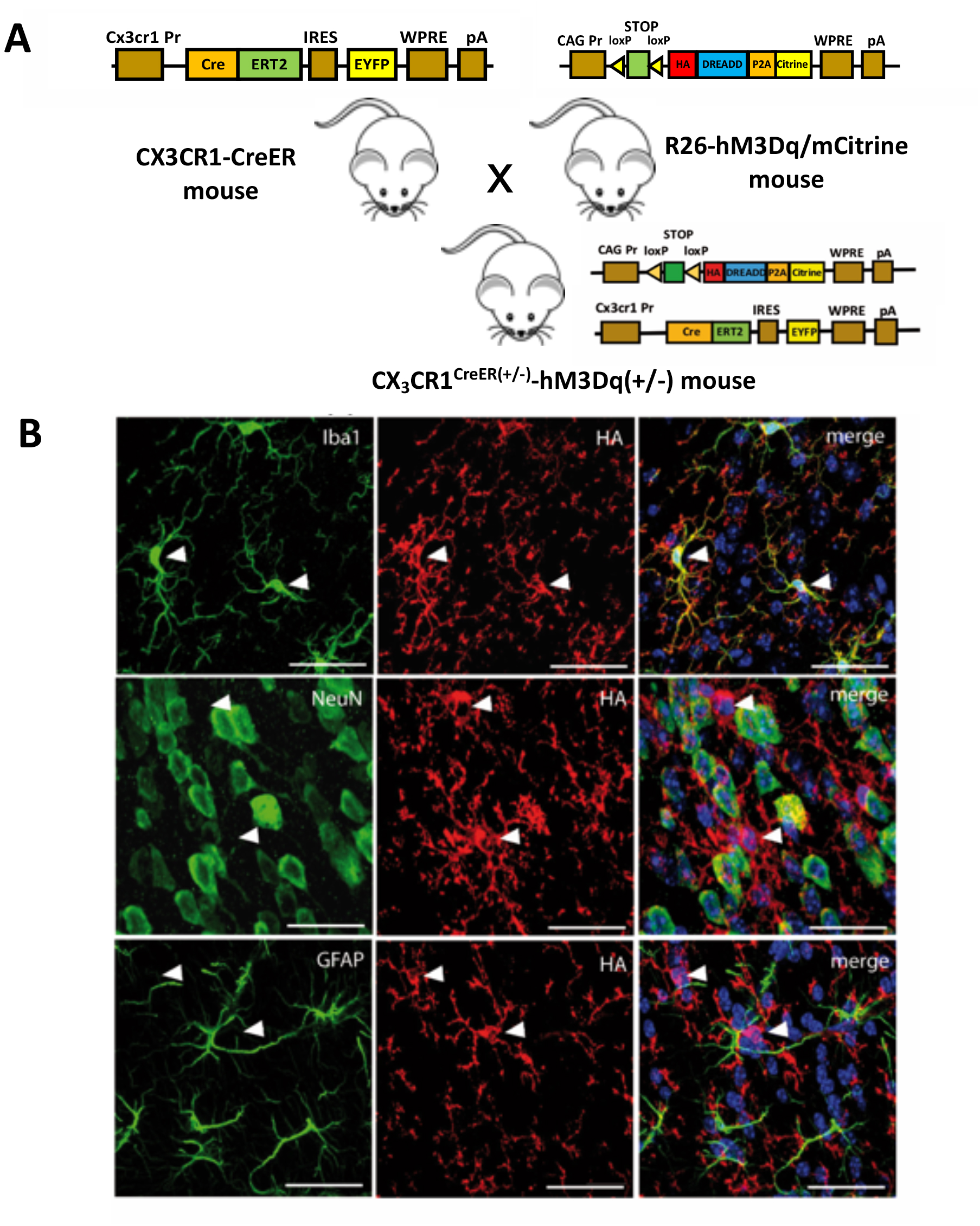
hM3Dq is expressed selectively in Iba1-positive microglia in CX^3^CR1^CreER(+/-)^-hM3Dq(+/-) mouse brain. **A)** Schematic of the targeting strategy used to obtain tamoxifen-inducible Cre-dependent expression of hM3Dq selectively in microglia. **B)** Representative confocal images of hippocampus of tamoxifen-treated CX^3^CR1*^CreER^*-hM3Dq(+/-) mice showing distribution of HA (red) and Iba1, NeuN or GFAP (green). Right panels show images of left panels merged with Hoechst nuclear stain (blue). Arrows indicate examples of HA-positive cells.

Brain sections co-stained for HA to label hM3Dq and markers of microglia (Iba1), neurons (NeuN) or astrocytes (GFAP, Fig. 1B) showed that hM3Dq was expressed mainly at the cell surface of Iba1-positive microglia, while virtually no co-localization with NeuN or GFAP was observed. A month after tamoxifen injection, 84.2% of counted cells were positive for both HA and Iba1, while 12.5% of cells were Iba1-positive only and 3.3% of cells were HA-positive only (n= 796 cells from three different mice). HA expression was not observed in either non-tamoxifen treated CX^3^CR1^CreER(+/-)^-hM3Dq(+/-) nor in CX^3^CR1^CreER(+/-)^ littermate control mice (data not shown).

### 3.2. Primary microglia cultures obtained from CX^3^CR1^CreER(+/-)^-hM3Dq(+/-) mice express functional DREADDs

To investigate the effect of hM3Dq activation on microglia, we initially isolated primary microglia from CX^3^CR1^CreER(+/-)^-hM3Dq(+/-) mice and treated cultures with the active tamoxifen metabolite 4-OHT to drive CreER-dependent hM3Dq expression (Fig. 2A). Immunofluorescence analysis showed colocalization of the microglia marker Iba1 with HA-tagged hM3Dq (Fig. 2B). HA expression was not detected in 4-OHT-treated control microglia cultures prepared from CX^3^CR1^CreER(+/-)^ littermates (Fig. 2B).

**Figure 2.**
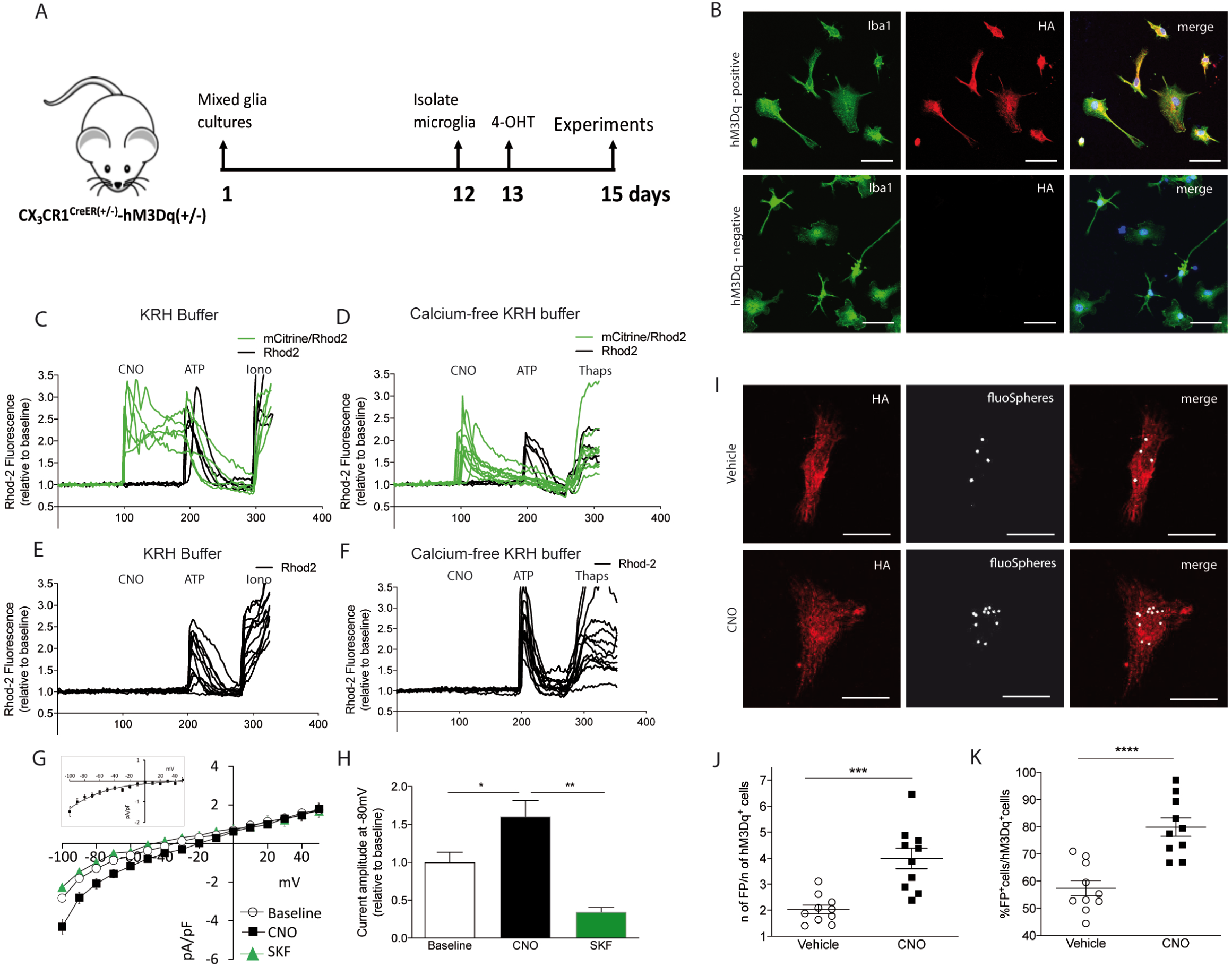
Microglia from CX^3^CR1^CreER^-hM3Dq(+/-) mice express functional hM3Dq upon tamoxifen treatment. **A)** Experimental approach and timeline of *in vitro* experiments. **B)** Representative images of immunostaining for Iba1 (green) and HA (red) in microglia culture treated with 4-OHT. **C-D)** Kinetics of relative Rhod-2 fluorescence in cells in response to 3 µM CNO, 1 mM ATP, and 2 µM ionomycin (iono) or 10 µM thapsigargin (thaps) in 4-OHT-treated cultures in KRH buffer (C) or calcium-free KRH buffer (D). **E-F)** Kinetics of relative Rhod-2 fluorescence in cells in non-treated cultures in KRH buffer (E) or calcium-free KRH buffer (F). **G)** Transmembrane current density (pA/pF) graph for CX^3^CR1*^CreER^*-hM3Dq(+/-) microglia in control conditions (baseline), and after being sequentially exposed to 3 µM CNO and 20 µM SKF-96365. Inset displays the CNO-evoked current obtained by baseline subtraction**. H)** Normalized current amplitude of CX^3^CR1^CreER^-hM3Dq(+/-) microglia (when V_M_ = −80mV) in conditions of control, and being exposed to CNO and SKF-96365 sequentially. **I)** Representative images of HA (red) showing phagocytosis of blue FluoSpheres (white) by vehicle-treated or CNO-treated CX^3^CR1^CreER^-hM3Dq(+/-) primary microglia. **J)** Quantification of the number of FluoSpheres per hM3Dq-positive microglia in cultures treated with vehicle or 5 µM CNO. **K)** Percentage of hM3Dq-positive microglia with FluoSpheres in cultures treated with vehicle or 5 µM CNO. Scale bar 20µm. *p<0.05, **p<0.01, ***p<0.001, ****p<0.0001. All data are represented as mean ± SEM.

Activation of h3MDq has been shown to increase intracellular Ca^2+^ signaling in cells (Pei et al., 2008; Urban and Roth, 2015). We used CNO (3 µM) to activate hM3Dq in Rhod2-loaded microglia isolated from CX^3^CR1^CreER(+/-)^-hM3Dq(+/-) mice and test for hM3Dq-mediated changes in intracellular Ca^2+^ signaling. hM3Dq–positive microglia cells were identified by the expression of mCitrine (Zhu et al., 2016). Application of CNO increased intracellular Ca^2+^ levels in the presence (Fig. 2C, green traces) and in the absence (Fig. 2D, green traces) of extracellular Ca^2+^ in hM3Dq–positive microglia cells, indicating mobilization from intracellular Ca^2+^ stores. Microglia isolated from CX^3^CR1^CreER(+/-)^ littermate control mice did not show a CNO-induced increase in Ca^2+^ signaling (Fig. 2C, D; Fig.2E, F, black traces). However, both positive and negative cells responded to ATP and ionomycin or thapsigargin, used as positive controls (Fig. 2C-F). These experiments indicate that hM3Dq expression in microglia allows selective activation of Ca^2+^ signaling pathways by CNO.

The release of Ca^2+^ from intracellular stores by hM3Dq is known to activate store-operated calcium channels (SOCC) that can replenish endoplasmic reticulum Ca^2+^ stores (Prakriya and Lewis, 2015). To test whether CNO-induced Ca^2+^ signaling activates SOC entry through a specific channel, we made voltage-clamp recordings in primary microglia from CX^3^CR1^CreER(+/-)^-hM3Dq(+/-) mice in a whole-cell configuration. By applying voltage ramps (from −100mV to +50mV over 150ms) to test cells, we found that activation of h3MDq by CNO evoked a current appearing only at negative membrane potentials (Figs. 2G and 2H; F_(2,36)_=7.663, p=0.0017, one-way ANOVA), which has been previously related to SOCCs (Michaelis et al., 2015). This CNO-evoked inward current in hM3Dq-expressing microglia was abolished in the presence of the SOCC inhibitor SKF-96365 (20 µM, Fig. 2G). To better demonstrate the current potentiated by CNO (p=0.0361, Tukey’s multiple comparisons test) and inhibited by SKF (p=0.0024), Fig. 2H shows the current at −80 mV of Fig 2G. Subtracting the control current from the CNO-evoked current displayed an inwardly rectifying current (Fig. 2G, inset) that is characteristic of store-operated Ca^2+^ release-activated Ca^2+^ (CRAC) currents (Ohana et al., 2009).

Elevation of intracellular Ca^2+^ has been shown to be important for many microglial functions including phagocytosis (Heo et al., 2015; Hoffmann et al., 2003; Toescu et al., 1998). Thus, we tested whether *in vitro* activation of hM3Dq by CNO facilitates phagocytosis (Fig. 2I-K). Treatment with CNO increased the number of FluoSpheres per hM3Dq-positive cell (Fig. 1J; t_(18)_=4.586, p=0.0002, Student’s t-test) as well as the percentage of hM3Dq-positive cells containing FluoSpheres (Fig. 2K; t_(18)_=5.133, p<0.0001, Student’s t-test) when compared to vehicle treatment. Taken together, these experiments show that microglia from CX^3^CR1^CreER(+/-)^-hM3Dq(+/-) mice express hM3Dq and that activation of this receptor by CNO triggers intracellular calcium increase and changes in microglial function.

### 3.3. Acute activation of hM3Dq increases mRNA expression of pro-inflammatory cytokines in the brain

As alterations in the intracellular calcium concentration have been shown to be important for modulation of cytokine expression by microglia (Goghari et al., 2000; Hoffmann et al., 2003; McLarnon et al., 2001; Tvrdik and Kalani, 2017), we investigated whether acute activation of hM3Dq alters microglia cytokine expression under normal conditions as well as during LPS-induced neuroinflammation. To test this, we used qPCR to measure mRNA expression of pro-inflammatory cytokines in the hippocampus of CX^3^CR1^CreER(+/-)^-hM3Dq(+/-) mice that were treated with CNO or vehicle prior to receiving either LPS or saline injection (Fig. 3A). Acute administration of CNO under non-inflammatory conditions (no LPS injection groups) increased mRNA levels of TNF-α (Fig. 3B; two-way ANOVA main effect of *CNO*: F_(1,26)_=101.5, p<0.0001, post-hoc Tukey’s analysis p<0.0001), IL-1β (Fig. 3C; two-way ANOVA main effect of *CNO*: F_(1,26)_=17.94, p<0.0001, post-hoc Tukey’s analysis p<0.0001) and IL-6 (Fig. 3D;, two-way ANOVA main effect of *CNO*: F_(1,26)_= 44.61, p<0.0001, post-hoc Tukey’s analysis p<0.0001). Although LPS treatment increased cytokine mRNA levels, a prior activation of hM3Dq by CNO (30 min before LPS injection, Fig. 3A) did not alter mRNA levels of TNF-α, IL-1β and IL-6 (Fig. 3B; TNF-α: two-way ANOVA LPS: F_(1,26)_= 44.22, p<0.0001, post-hoc Tukey’s analysis p=0.21; Fig. 3C; IL-1β: LPS: F_(1,26)_= 50.97, p<0.0001, post-hoc Tukey’s analysis p=0.96; and Fig. 3D; IL-6: two-way ANOVA LPS: F_(1,26)_= 69.62, p<0.0001, post-hoc Tukey’s analysis p=0.54), possibly due to ceiling effects.

**Figure 3.**
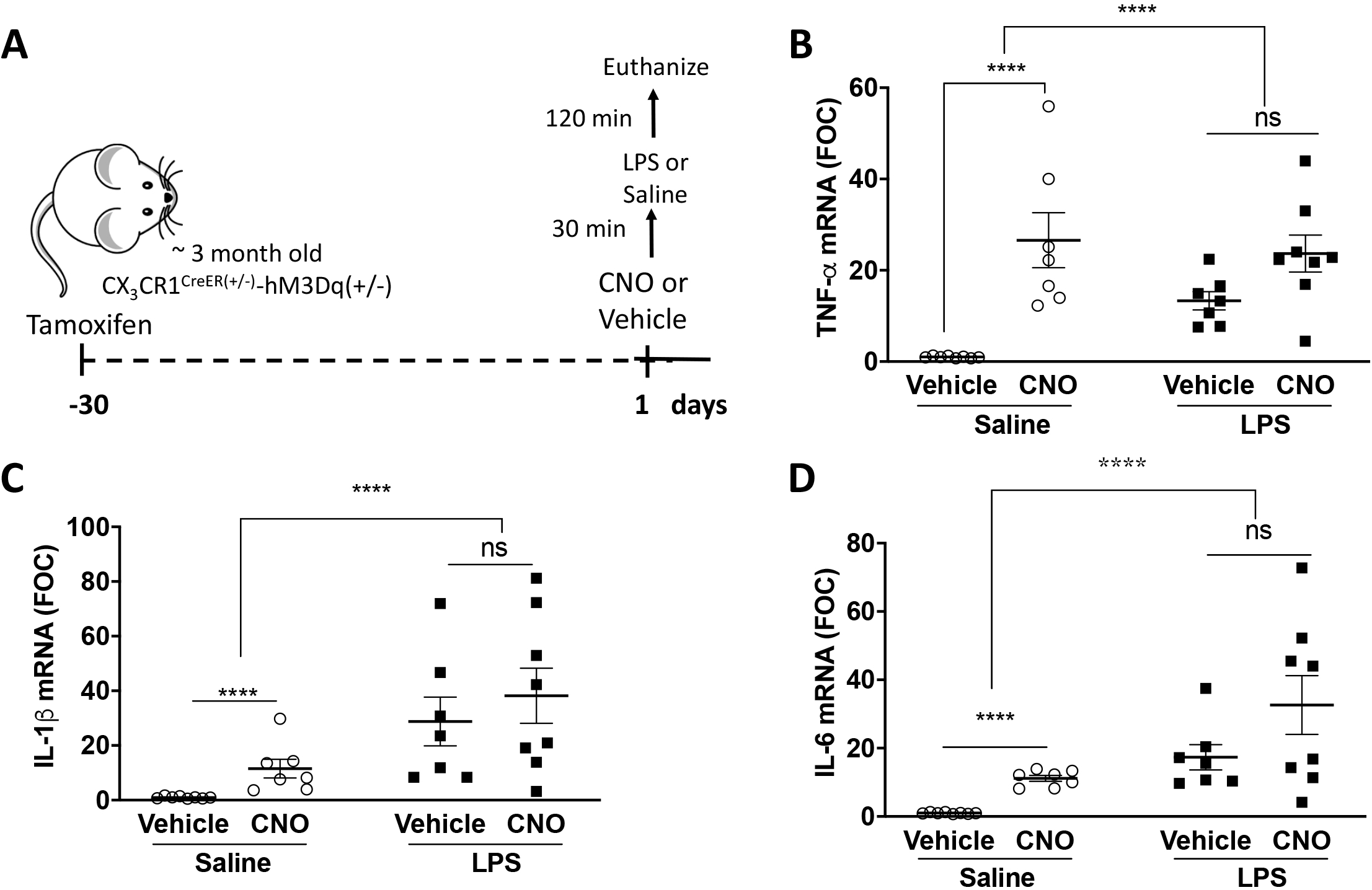
Acute microglial hM3Dq activation by CNO increases cytokine expression in the brain. **A)** Experimental approach and timeline for hippocampus gene expression experiments. CNO (1 mg/kg; i.p.); LPS (0.1 mg/kg; i.p.). **B-D)** qPCR measurements of mRNA expression of TNF-α (B), IL-1β (C), and IL-6 (D) in hippocampus of hM3Dq-positive mice following vehicle or CNO treatment prior to saline or LPS injection. All data are expressed as folds of change from the vehicle/saline-treated group and are represented as mean ± SEM. ns p>0.05, ****p<0.0001.

### 3.4. Activation of hM3Dq in microglia does not cause overt behavioral change

Activation of pro-inflammatory cytokine expression in the brain by systemic inflammatory challenges induces symptoms of sickness behavior with a number of behavioral modifications including decreased locomotion, decreased exploratory behavior, increased anxiety, anorexia, and weight loss (Dantzer, 2001; Tchessalova et al., 2018). To examine whether increased expression of pro-inflammatory cytokines by hM3Dq activation in microglia also leads to behavioral changes we conducted a series of behavioral analyses where over 3 days mice were treated with CNO (1.0 mg/kg; i.p.) or vehicle (0.5% DMSO in saline) 30 min prior to individual behavioral tests (Fig. 4A).

**Figure 4.**
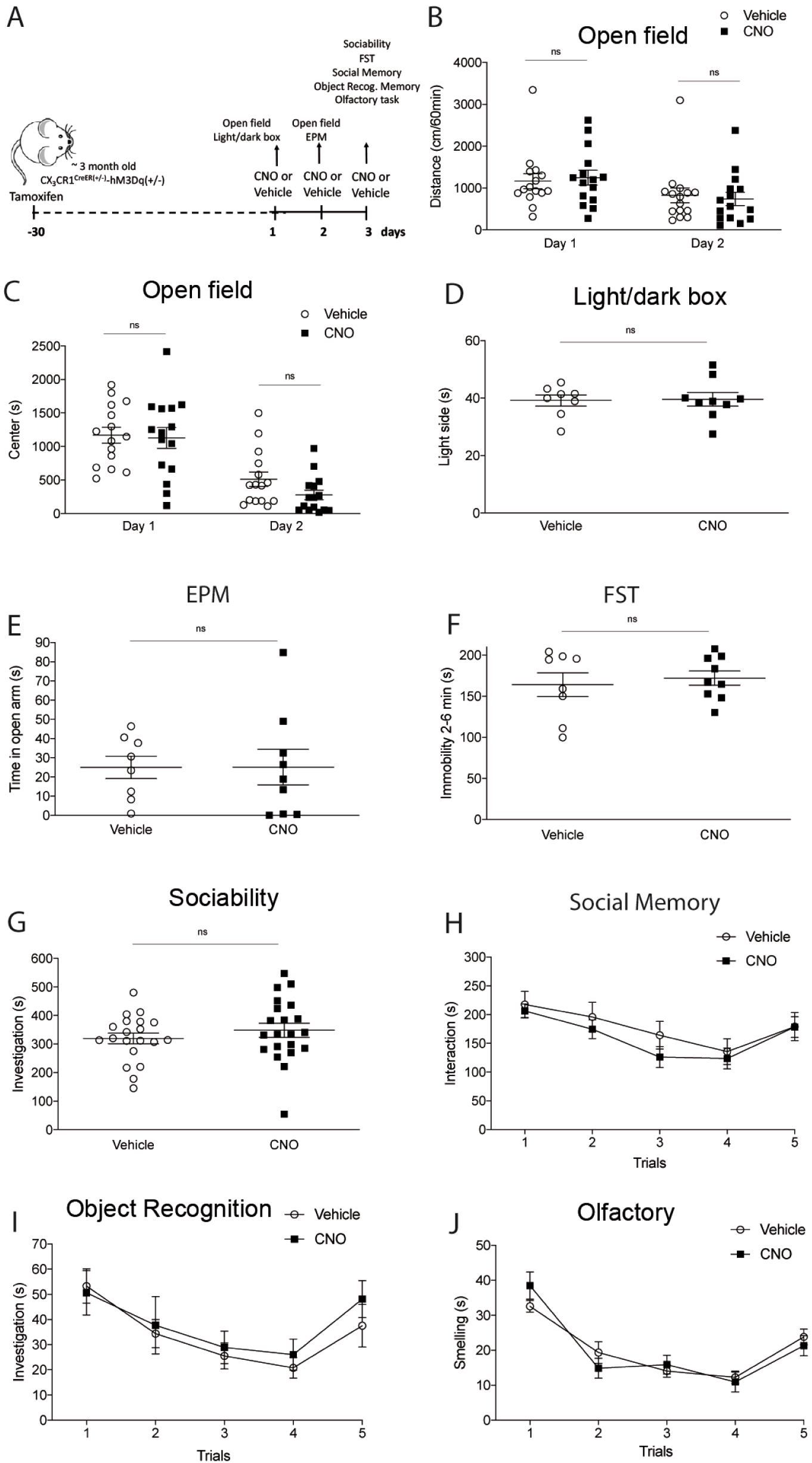
Microglial hM3Dq activation by CNO does not cause overt behavioral change. **A)** Timeline of CNO treatment and behavior assessments. CNO (1.0 mg/kg; i.p.) or vehicle was injected 30 min before each behavioral task. **B-C**) Total distance travelled (B) and time spent in the center (C) of an open field on two consecutive days. **D)** Time spent in the light side of a light-dark box. **E)** Time spent in the open arm of an elevated plus maze (EPM). **F)** Time spent immobile during minutes 2-6 in forced swim test (FST). **G)** Time spent investigating a juvenile mouse in a three-chamber social preference task (Sociability). **H)** Time spent interacting with an intruder mouse for social memory assessment. *1 trial = 5 min*. **I)** Time spent investigating a novel object to assess object recognition. **J)** Time spent investigating an enclosed scent to test olfaction. Data are represented as mean ± SEM. ns p>0.05.

Investigation of spontaneous activity in the open field using automated locomotor boxes showed no significant difference between CX^3^CR1^CreER(+/-)^-hM3Dq(+/-) mice treated with CNO or vehicle on the distance travelled in 60 min, both on day 1 and day 2, indicating that hM3Dq activation does not interfere with locomotor activity (Fig. 4B; no effect of CNO on distance travelled (*F*_(1,28)_=3.06, p=0.995); no interaction between CNO treatment and time (*F*_(1,28)_= 0.237, p=0.630). Additionally, both CNO- and vehicle-treated CX^3^CR1^CreER(+/-)^-hM3Dq(+/-) mice showed the expected decrease in locomotor activity on the 2^nd^ test day due to habituation (Fig. 4B; significant effects of day on the total distance travelled; *F*_(1,28)_=5.947, p=0.0213, two-way ANOVA repeated measures).

We tested for changes in anxiety-like behavior using different paradigms. Time spent in the center vs. the periphery of the open field (Fig. 4C) was not affected by activation of hM3Dq (No main effect of *CNO treatment*: *F*_(1,28)_=1.045, p=0.3153; No treatment x day *interaction*: *F*_(1,28)_=1.031, p=0.3187, two-way ANOVA repeated measures). Likewise, we found that activation of hM3Dq with CNO did not affect the time mice spent exploring a novel unprotected environment in the light-dark box (light compartment) (Fig. 4D; t_(15)_=0.14, p=0.8907, Student’s t-test) or the elevated plus maze (open arms) (Fig. 4E; t_(15)_=0.0075, p=0.9941, Student’s t-test). Additionally, we did not detect any effect of hM3Dq activation in the forced swim test (Fig. 4F; t_(15)_=0.48, p=0.6391, Student’s t-test).

We also investigated whether activation of hM3Dq with CNO alters social behaviors. The three-chamber social preference task showed that hM3Dq signaling in microglia does not change sociability, as time spent investigating another rodent as compared to time spent investigating a plastic block was similar between CNO- and vehicle-treated CX^3^CR1^CreER(+/-)^-hM3Dq(+/-) mice (Fig. 4G; t_(39)_=0.93, p=0.359, Student’s t-test). To investigate social memory, we tested mice on a habituation-dishabituation paradigm using a mouse intruder. CNO- and vehicle-treated CX^3^CR1^CreER(+/-)^-hM3Dq(+/-) mice showed significantly more exploration of the intruder during first contact and decreased exploration time with subsequent exposure to the same juvenile, suggesting that both cohorts could equally remember the initial intruder. Upon changing to an unfamiliar mouse (fifth trial), both CNO- and vehicle-treated-mice explored the new intruder as much as they explored the original intruder during the first contact, confirming their preference for social novelty (Fig. 4H; No effect of *CNO treatment*: *F*_(1,28)_=0.399, p=0.533; a significant effect of trial on time of *interaction: F*_(4,112)_=21.84, p<0.0001, two-way ANOVA repeated measures). A similar paradigm was used to investigate object and olfactory memory and, likewise, no difference between CNO- and vehicle-treated CX^3^CR1^CreER(+/-)^-hM3Dq(+/-) mice was observed (Fig. 4I; object recognition memory: two-way ANOVA repeated measures no main effect of *CNO treatment*: *F*_(1,13)_=0.1905, p=0.6697; no treatment x trial *interaction*: *F*_(4,52)_= 0.8272, p=0.5139; Fig. 4J; olfactory memory: no main effect of *CNO treatment*: *F*_(1,14)_=0.004324, p=0.9485; no treatment x trial *interaction*: *F*_(4,56)_=1.156, p=0.3403). These results suggest that, within the parameters we used, activation of hM3Dq in microglia does not affect normal mouse behavior.

### 3.5. Chronic stimulation of hM3Dq in microglia attenuates LPS-induced increase of pro-inflammatory cytokines in the brain

Repetitive peripheral administration of LPS has been shown to induce immune memory in microglia that can then modify their response to inflammatory stimuli (Wendeln et al., 2018). To test the possibility that repetitive activation of hM3Dq could affect the subsequent microglial responses to LPS, we treated CX^3^CR1^CreER(+/-)^-hM3Dq(+/-) mice for 4 days with vehicle or CNO (1 mg/kg), then challenged them with LPS 2 hours before collecting brain tissues for evaluation of mRNA expression of the pro-inflammatory cytokines TNF-α, IL-1β, or IL-6 (Fig. 5A). Interestingly, with the 4-days CNO treatment, the chronic activation of microglial hM3Dq alone no longer increased mRNA for inflammatory cytokine (TNF-α: Fig. 5B, two-way ANOVA no main effect of *CNO*: F_(1,37)_=3.86, p=0.057; post-hoc Tukey’s analysis; IL-1β: Fig. 5C, two-way ANOVA no main effect of *CNO*: F_(1,37)_=2.69, p=0.11; post-hoc Tukey’s analysis; and IL-6: Fig. 5D, two-way ANOVA no main effect of *CNO*: F_(1,38)_= 1.89, p=0.18; post-hoc Tukey’s analysis). Remarkably, in LPS-treated mice, chronic activation of hM3Dq with CNO significantly attenuated LPS-induced mRNA levels of TNF-α (Fig. 5B; two-way ANOVA main effect of *LPS*: F_(1,37)_=18.14, p=0.0001; post-hoc Tukey’s analysis Vehicle-LPS x CNO-LPS p=0.02), IL-1β (Fig. 5C; two-way ANOVA main effect of *LPS*: F_(1,37)_=74.53, p<0001; post-hoc Tukey’s analysis Vehicle-LPS x CNO-LPS p=0.02), and IL-6 (Fig. 5D; two-way ANOVA main effect of *LPS*: F_(1,38)_=46.67, p<0.0001; post-hoc Tukey’s analysis Vehicle-LPS x CNO-LPS p=0.047). In littermate control mice that do not express hM3Dq (CX^3^CR1^CreER(+/-)^) and were submitted to the same protocol CNO had no effect (Fig. 5E-G), validating our findings that modulation of LPS-induced cytokine mRNA was attributable to hM3Gq signaling.

**Figure 5.**
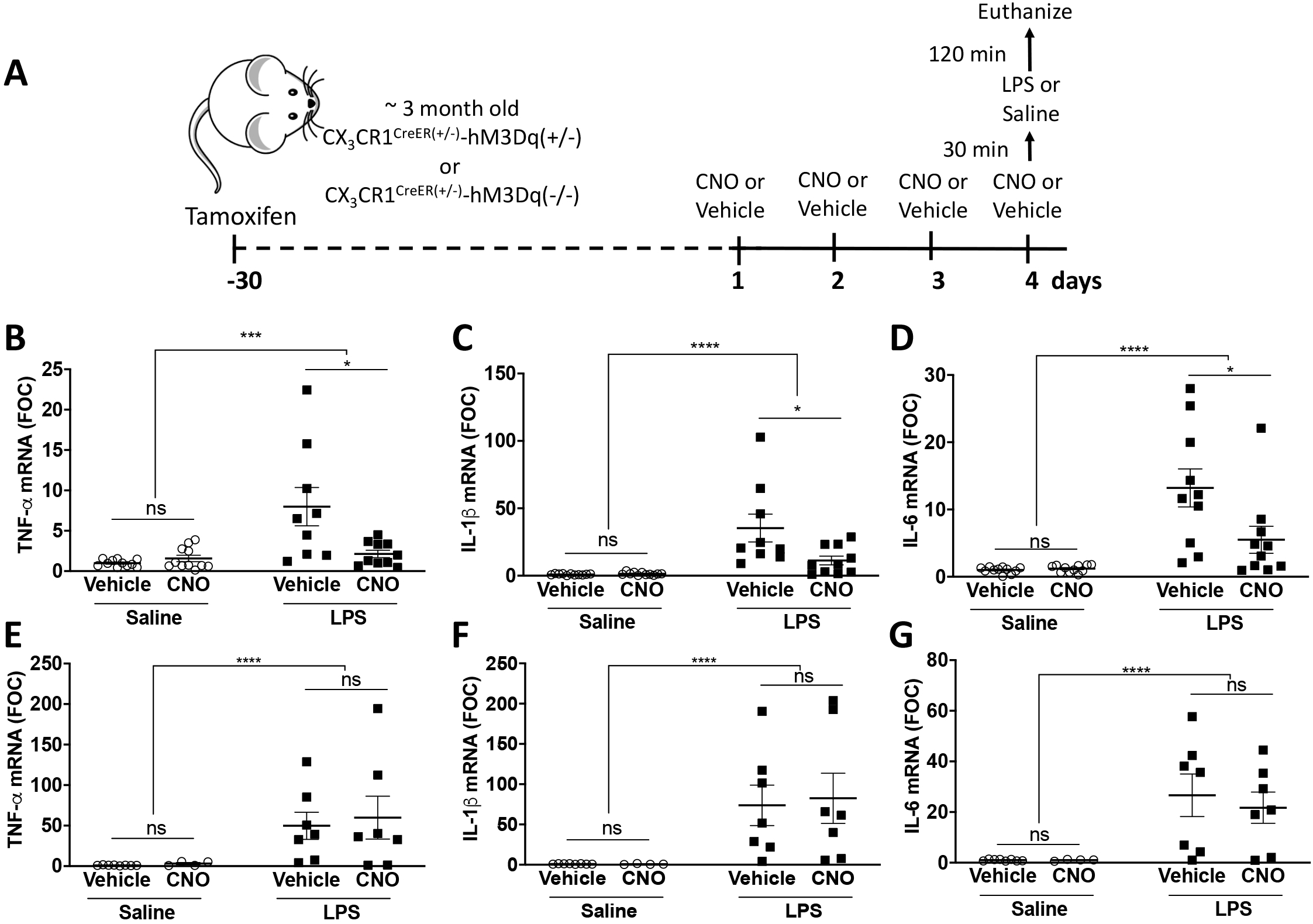
Chronic microglial hM3Dq activation by CNO attenuates LPS-induced increase of pro-inflammatory cytokines in the brain. **A)** Experimental approach and timeline for hippocampus gene expression experiments. CNO (1 mg/kg; i.p.); LPS (0.1 mg/kg; i.p.). **B-D)** qPCR measurements of mRNA expression of TNF-α (B), IL-1β (C), and IL-6 (D) in hippocampus of CX^3^CR1^CreER(+/-)^-hM3Dq(+/-) mice 2 hr after LPS or saline injection. **E-G)** qPCR measurements of mRNA expression of TNF-α (E), IL-1β (F), and IL-6 (G) in hippocampus of CX^3^CR1^CreER(+/-)^ (hM3Dq-negative) mice. All data are expressed as folds of change from the vehicle/saline-treated group and are represented as mean ± SEM. ns p>0.05, *p<0.05, *\*\*\* p<0.001, **** p<0.0001*.

### 3.6. Chronic stimulation of hM3Dq in microglia prevents LPS-induced sickness-like behavior

To examine whether prior hM3Dq activation in microglia influences LPS-induced behavioral changes, we treated CX^3^CR1^CreER(+/-)^-hM3Dq(+/-) mice with CNO for 4 consecutive days, challenged them with LPS (0.1 mg/kg, i.p.) on the 4^th^ day and then performed different behavioral tests 2 h and 24 h after LPS administration (Fig. 6A). As control groups we used CX^3^CR1^CreER(+/-)^-hM3Dq(+/-) mice that received vehicle instead of CNO. Weight analysis during the 5 days showed that activation of hM3Dq with CNO alone did not lead to significant change in weight. On the other hand, both CNO- and vehicle-treated mice showed significant weight loss after LPS injection (Fig. 6B; *time*: F_(4,80)_=51.43, p<0.0001, two-way ANOVA repeated measures).

**Figure 6.**
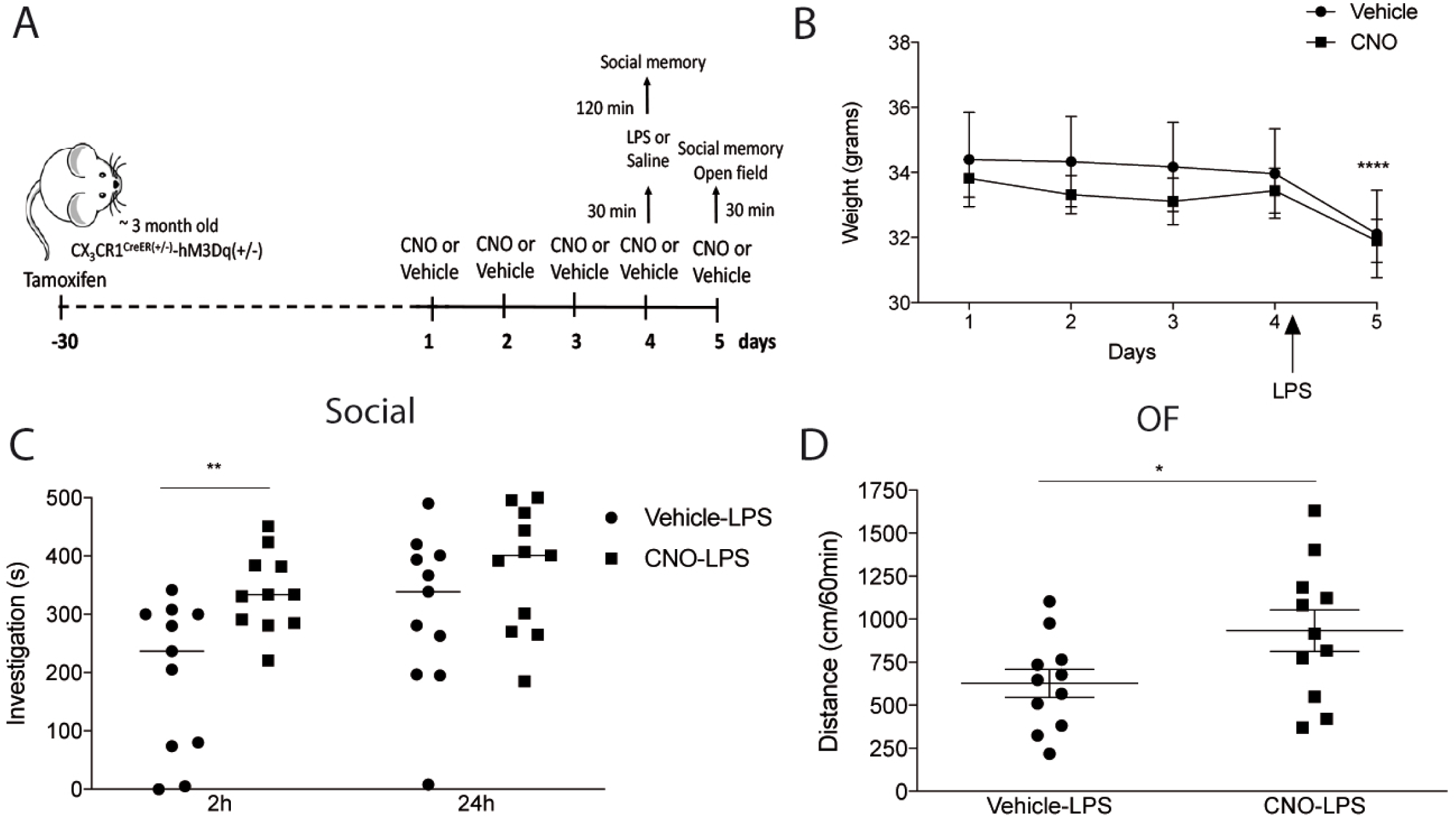
Chronic stimulation of hM3Dq in microglia prevents LPS-induced sickness behavior. **A)** Experimental approach and timeline for CNO treatment, LPS challenge and behavior assessments. CNO (1 mg/kg; i.p.); LPS (0.1 mg/kg; i.p.). **B)** Body weight decreased after LPS injection (day 4) for both CNO- and vehicle-treated mice (*n=11 per treatment, **** in comparison to days 1-4*). **C)** Time spent investigating juvenile mouse in a three-chamber social preference task at 2h and 24h after LPS treatment. **D)** Locomotor activity in an open field 24h after LPS treatment. All data are represented as mean ± SEM. *\* p<0.05, ** p<0.01, **** p<0.0001*.

Mice in which hM3Dq was activated by CNO showed longer social interaction in the three-chamber social preference task than vehicle-treated mice when examined at 2 hour after LPS injection (Fig. 6C; main effect of CNO *treatment*: F_(1,20)_=8.946, p=0.0072, main effect of time *time*: F_(1,20)_=5.46, p=0.029, *interaction*: *F*_(1,20)_=1.318, p= 0.2646, two-way ANOVA). Comparison of social interaction data from Figures 4G and 6C shows that vehicle-treated CX^3^CR1^CreER(+/-)^-hM3Dq(+/-) mice that did not receive LPS explored the juvenile mice much longer (319.8 ±18.37; Fig 4G; N=20) than vehicle-treated mice injected with LPS (193.72 ± 39.02; Fig. 6C; N=11), indicating that, as expected (Corona et al., 2010), peripheral LPS injection decreased social interaction. Importantly, exploratory activity of CNO-treated mice injected with LPS (338.00 ± 20.51; Fig 6C; N=11) was very similar to that of mice that did not receive LPS, with or without CNO (319.8 ± 18.37; Fig 4G; N=20). These results suggest that chronic activation of hM3Dq in microglia protects against LPS-induced suppression of social interaction. Twenty-four hours after LPS injection, vehicle-injected mice showed increased social interaction (Fig. 6C), consistent with the time course of recovery of social interaction following low-dose LPS injection (Corona et al., 2010). At the 24h time point, there was no difference between vehicle- and CNO-treated groups. A similar protective effect was observed when we tested mice for locomotor activity in the open field 24h after LPS injection (compare Fig. 6D and 4B). Together, these data suggest that chronic activation of hM3Dq in microglia protects against LPS-induced sickness behavior.

## Discussion

Using the Cre-lox system to drive specific and inducible expression of hM3Dq in microglia, we dissected the action of muscarinic M3 receptor (the hM3Dq progenitor) signaling in these cells *in vivo*, with implications for other GPCRs that are also Gq-associated. Our results demonstrate that acute activation of microglial hM3Dq-signaling enhances the production of pro-inflammatory cytokines in the brain. In contrast, chronic activation of this pathway attenuates LPS-mediated cytokine expression and sickness behavior.

Tamoxifen-induced Cre expression in CX^3^CR1-positive cells allowed persistent expression of hM3Dq in long-lived microglia but not in short-lived myeloid cells. Previous validation of the CX^3^CR1^CreER^-dependent manipulation of gene expression in microglia using the reporter Rosa26-STOP-DsRed mice showed that in the absence of tamoxifen, very few (∼0.3%) CX^3^CR1-positive cells expressed DsRed. In contrast, 5 days after tamoxifen treatment, the majority of CX^3^CR1-positive cells were positive for DsRed. Also, 30 days following tamoxifen treatment, more than 90% of microglia still expressed DsRed (Parkhurst et al., 2013), whereas less than 2% of CX^3^CR1-positive monocytes in the blood showed the expression due to their rapid turnover. Importantly, because microglia are long-lived (Fuger et al., 2017), they continue to express the recombined gene (Parkhurst et al., 2013). Our results are compatible with these previous observations, as we detected hM3Dq in microglia up to almost a year after tamoxifen treatment (not shown). Also, we did not detect expression of hM3Dq in astrocytes or neurons. These results confirm that the mouse model we generated expresses hM3Dq specifically in microglia.

Microglia express several GPCRs that associate to Gq-signaling heterotrimeric G proteins, such as group 1 metabotropic glutamate receptors (mGluR1, mGluR5) (Biber et al., 1999), purinergic receptors (P2Y1, P2Y2, P2Y4, P2Y6, P2Y11) (Koizumi et al., 2013), adrenergic (α1AR) (Mori et al., 2002) and the M3 muscarinic receptor (Pannell et al., 2016). Although these receptors are activated by different ligands, they are all able to elicit Gq pathway activation. Gq proteins are known to mobilize calcium through inositol 1,4,5-triphosphate-regulation of intracellular stores and diacylglycerol-dependent protein kinase C activation (Mizuno and Itoh, 2009). Our *in vitro* experiments showed that CNO was able to elicit an increase in microglial intracellular calcium concentration, only in hM3Dq-positive cells. This increase was seen in both calcium-free media and media containing calcium, indicating that hM3Dq was functional and the source of calcium influx is, at least in part, from intracellular stores. The functionality of hM3Dq and the signaling pathway activated were also confirmed by patch-clamp experiments showing that hM3Dq activation by CNO induced an increase in conductance in microglia that was abolished by the SOCC inhibitor.

Our results also showed that activation of hM3Dq by CNO increased phagocytosis in cultured microglia. Conversely, activation of muscarinic receptors by carbachol led to increased chemotaxis and decreased phagocytosis in microglia (Pannell et al., 2016). It is possible that the difference in results stems from the fact that while CNO allows for specific activation of M3 muscarinic receptors, carbachol, a non-selective cholinergic agonist, activates all subtypes of muscarinic and also nicotinic receptors. To note, activation of P2Y6 receptor, a Gq-linked GPCR that responds to UDP, also increased microglia phagocytosis (Koizumi et al., 2007; Neher et al., 2014).

Microglia sense exogenous and endogenous inflammatory signals and orchestrate subsequent neuroinflammation which triggers a number of behavioral changes (Biesmans et al., 2013). Accumulating evidence indicate that the microglial response to inflammatory stimuli can be very plastic (Neher and Cunningham, 2019). Upon previously encountering an inflammatory stimulus, microglia can adjust their activation state and elicit amplified responses to a secondary inflammatory insult, a phenomenon referred to as microglial pre-conditioning or “priming” (Neher and Cunningham, 2019). Microglia can also develop long-lasting molecular reprograming that suppresses microglial activation upon a secondary insult, a phenotype referred to as “innate immune memory” or tolerance (Netea et al., 2016; Netea et al., 2011). Microglia have recently been shown to display immune training and tolerance in response to repeated daily peripheral stimulation with LPS. One low-dose intraperitoneal injection of LPS led to increased pro-inflammatory cytokine levels in the brain in response to the second injection. Conversely, upon the fourth daily LPS injection microglia barely responded, showing decreased cytokine levels in the brain, an indication of immune tolerance (Wendeln et al., 2018). We showed that acute activation of hM3Dq in microglia strongly induced brain TNF-α, IL-1β and IL-6 mRNA expression, however, this “pro-inflammatory” effect was completely lost after repeated (4 days) hM3Dq activation, suggesting that repeated activation of muscarinic M3 receptors in microglia can result in microglial tolerance. Recurrent hM3Dq activation also diminished LPS-induced cytokine production and prevented LPS-induced sickness behavior, raising an intriguing possibility that hM3Dq-signaling alone could lead to microglial immune memory formation. Remarkably, pharmacological studies using muscarinic M3-knockout mice showed that in the periphery, M3-signaling is important for immune memory formation (Darby et al., 2015). M3-knockout mice showed reduced CD4+ T cell activation and cytokine production, which resulted in impaired ability to resolve a secondary infection (Darby et al., 2015).

It is important to note that neither chronic nor acute activation of hM3Dq in microglia affected normal mouse behavior, indicating that hM3Dq activation per se does not lead to sickness behavior or other deleterious effects. Thus, these results suggest that chemogenetics can be a powerful approach to modulate microglia and modify neuroinflammation.

Activation of microglia using DREADDs has been reported previously using viral constructs (Grace et al., 2016; Grace et al., 2018). In these studies, hM3Dq or hM4Di DREADDs were expressed in spinal cord microglia by intrathecal administration of AAV9 vector constructs. We avoided using this approach because microglia can respond to viral infection (Kunitsyna et al., 2016; Martianova et al., 2019), which may be a confounder in these types of studies. In virus-infected animals, CNO-induced activation of hM3Dq was able to elicit allodynia in male rats (Grace et al., 2018). In the present study, CX^3^CR1^CreER(+/-)^-hM3Dq(+/-) mice showed no visual signs of pain following either chronic or acute CNO treatment, based on the basal behavioral performance that was not different from controls. However, because we did not specifically test allodynia, we cannot compare our observations to their results. In agreement with our observations with acute CNO treatment, Grace and colleagues also showed increased pro-inflammatory cytokine synthesis when activating hM3Dq signaling acutely in BV-2 microglia cells.

Our data suggest that activation of the muscarinic Gq pathway could be involved in inflammatory cytokine production and immune memory formation, which can likely be extended to other Gq-activating GPCRs. We provide a proof of principle that the use of chemogenetics *in vivo* can be a powerful approach to modulate microglia and regulate cytokine production and the resulting sickness behavior. This approach allows the manipulation of microglial activity *in vivo* and the ability to study the cross-talk between microglia and neurons in different settings, including during development and in mouse models of neurodegenerative diseases.

## Acknowledgements

hM3Dq mutant mice were a generous gift of Ute Hochgeschwender and Bryan Roth. We thank Matt Cowan, Jue Fan and Sanda Raulic for assistance with mice and behavioral experiments.

